# Zika virus infection induces IL-1β-mediated inflammatory responses by macrophages in the brain of an adult mouse model

**DOI:** 10.1101/2023.01.05.522967

**Authors:** Gi Uk Jeong, Sumin Lee, Do Yeon Kim, Jaemyun Lyu, Gun Young Yun, Junsu Ko, Young-Chan Kwon

**Affiliations:** Department of Convergent Research for Emerging Virus Infection, Korea Research Institute of Chemical Technology, Daejeon 34114, Republic of Korea; Arontier Co., Ltd., Seoul 06735, Republic of Korea

**Author notes:** Address correspondence to Young-Chan Kwon. Gi Uk Jeong and Sumin Lee contributed equally to this work. Sumin Lee, International Vaccine Institute, Seoul, Republic of Korea.

**Keywords:** Zika virus, IL-1β, macrophage, neuroinflammation, Complement C3

## Abstract

During the 2015/16 Zika virus (ZIKV) epidemic, ZIKV associated neurological diseases were reported in adults, including microcephaly, Guillain-Barre syndrome, myelitis, meningoencephalitis, and fatal encephalitis. However, the mechanisms underlying the neuropathogenesis of ZIKV infection are not yet fully understood. In this study, we used an adult ZIKV-infection mouse model (*Ifnar1*^−/−^) to investigate the mechanisms underlying neuroinflammation and neuropathogenesis. ZIKV infection induced the expression of proinflammatory cytokines, including IL-1β, IL-6, IFN-γ, and TNF-α, in the brains of *Ifnar1*^−/−^ mice. RNA-seq analysis of the infected mouse brain also revealed that genes involved in innate immune responses and cytokine-mediated signaling pathways were significantly upregulated at 6 days post infection. Furthermore, ZIKV infection induced macrophage infiltration and activation, and augmented IL-1β expression, whereas microgliosis was not observed in the brain. Using human monocyte THP-1 cells, we confirmed that ZIKV infection promotes inflammatory cell death and increases IL-1β secretion. In addition, the expression of complement component C3, which is associated with neurodegenerative diseases and known to be upregulated by proinflammatory cytokines, was induced by ZIKV infection through the IL-1β-mediated pathway. An increase in C5a produced by complement activation in the brains of ZIKV-infected mice was also confirmed. Taken together, our results suggest that ZIKV infection of the brain in this animal model augments IL-1β expression in infiltrating macrophages and elicits IL-1β-mediated inflammation, which can lead to the destructive consequences of neuroinflammation.

**Importance:** Zika virus (ZIKV) associated neurological impairments are an important global health problem. Our results suggest that ZIKV infection of the mouse brain can induce IL-1β-mediated inflammation and complement activation, contributing to the development of neurological disorders. Thus, our findings reveal a mechanism by which ZIKV induces neuroinflammation in the mouse brain. Although we used adult type I IFN receptor IFNAR knockout (*Ifnar1*^−/−^) mice owing to the limited mouse model of ZIKV pathogenesis, our conclusion could contribute to understanding ZIKV associated neurological diseases to develop treatment strategies based on these findings for the patients with ZIKV infection.

## Introduction

During the 2015/16 epidemic of Zika virus (ZIKV) in the Americas (1), case reports indicated an association between congenital ZIKV infection and microcephaly (2, 3). The rapid spread of ZIKV has also indicated a causal relationship between ZIKV infection and neurological complications in adults, including Guillain-Barré syndrome (GBS) (4), myelitis (5), meningoencephalitis (6), and fatal encephalitis (7), suggesting a possible association between ZIKV and neurological diseases. Limb weakness, hyporeflexia/areflexia, facial palsy, and paresthesia are the most frequent neurological symptoms in patients with ZIKV-associated GBS (8). However, the neuropathogenesis of ZIKV infection is not yet fully understood.

ZIKV belongs to the *Flaviviridae* family of RNA viruses and is a neurotropic flavivirus, along with Japanese encephalitis and West Nile viruses (9). The detection of ZIKV RNA in the brains and cerebrospinal fluid of adult patients with ZIKV-induced neurological disorders also suggests a neuroinvasive characteristic of ZIKV (6, 10-13). ZIKV can infect human neuronal progenitor cells (NPCs), leading to cell death and abnormal growth (14, 15). ZIKV infection of NPCs in the adult mouse brain results in cell death and reduced proliferation (16). In addition to NPCs, human placental macrophages (17) and glial cells (18) are also susceptible to ZIKV infection. Lum et al. demonstrated that ZIKV infects human fetal brain microglia and macrophages, which induces high levels of proinflammatory cytokines; these are implicated in ZIKV-derived neuroinflammation (19). Increasing evidence indicates that neuroinflammation is a major contributor to the pathogenesis of neurological diseases (20). Thus, an investigation to address the mechanisms underlying the neuropathogenesis of ZIKV infection would help to understand ZIKV-associated neurological diseases.

Proinflammatory cytokines are central mediators of the inflammatory response. Interleukin (IL)-1β is one of the most extensively studied cytokines involved in neuroinflammation and neurodegenerative diseases (21). In the brains of patients with human immunodeficiency virus 1 encephalitis, IL-1β expression is increased in infiltrating macrophages, microglia, and astrocytes (22, 23). Complement component 3 (C3) is also induced in neurodegenerative diseases and is upregulated by proinflammatory cytokines such as IL-1β, interferon (INF)-γ, and tumor necrosis factor (TNF)-α (24-27). Simian immunodeficiency virus infection of the central nervous system (CNS) in rhesus macaques induces C3 expression in infiltrating macrophages, astrocytes, and neurons (28). Although both IL-1β and C3 can play neuroprotective roles in immune responses, uncontrolled biosynthesis and activation can lead to critical brain tissue damage (24, 29). Hence, whether IL-1β and C3 expression in the brain is affected by ZIKV infection needs to be investigated to explore ZIKV-associated neurological diseases.

Immune-competent adult mice are resistant to ZIKV infection in part because ZIKV fails to effectively antagonize Stat2-dependent IFN responses in mice despite ZIKV NS5 protein binds and degrades STAT2 for the immune evasion. (30-33). Thus, we used adult type I IFN receptor IFNAR knockout (*Ifnar1*^−/−^) mice as a ZIKV infection mouse model to examine proinflammatory responses in the CNS and the neuropathogenesis of ZIKV infection. RNA-seq analysis was performed to examine neuroinflammation in response to ZIKV infection in the brains of *Ifnar1*^−/−^ mice. We focused on the immune cells in the central nervous system that are susceptible to ZIKV infection and consequently contribute to neuroinflammation in this animal model.

## Results

### ZIKV infection of the brain induces proinflammatory responses in *Ifnar1*^−/−^ mice

When *Ifnar1*^−/−^ C57BL/6 mice were infected with 10^3^ plaque-forming units (PFU) of the PRVABC59 strain of ZIKV via a subcutaneous route, most infected mice became paralyzed on at least one hind limb at 9 days post infection (dpi) (Fig. 1A). To our knowledge, there are two mechanisms underlying ZIKV-associated paralysis in *Ifnar1*^−/−^ mice. First, ZIKV infection of astrocytes breaks down the blood-brain barrier of *Ifnar1*^−/−^ mice, leading to a large influx of CD8+ T cells that promotes paralysis (34). Second, ZIKV infection in the spinal cord of *Ifnar1*^−/−^ mice results in motor neuron synaptic retraction and inflammation (35). To address whether infection in the brain or spinal cord accounts for paralysis in ZIKV infected mice, we assessed the viral load at 6 dpi in the brain and spinal cord along with that in the eyes, kidneys, testes, and ovaries, which are reported as ZIKV susceptible organs (Fig. 1B). High levels of viral RNA were detected in both the brain and spinal cord, but the levels in the spinal cord were higher than those in the brain. Next, we analyzed the mRNA levels of proinflammatory cytokines, including IL-1β, IL-6, IFN-γ, and TNF-α, in the brain and spinal cord using RT-qPCR at 6 dpi (Fig. 1C). Notably, the mRNA expression of proinflammatory cytokines in the brains of ZIKV-infected mice was much higher than that in the spinal cord. Upregulated proinflammatory cytokine expression in the ZIKV-infected mouse brain was confirmed using ELISA (Fig. 1D). These results indicated that ZIKV infection of the brain induces proinflammatory responses that contribute to neuroinflammation.

**Fig. 1.**
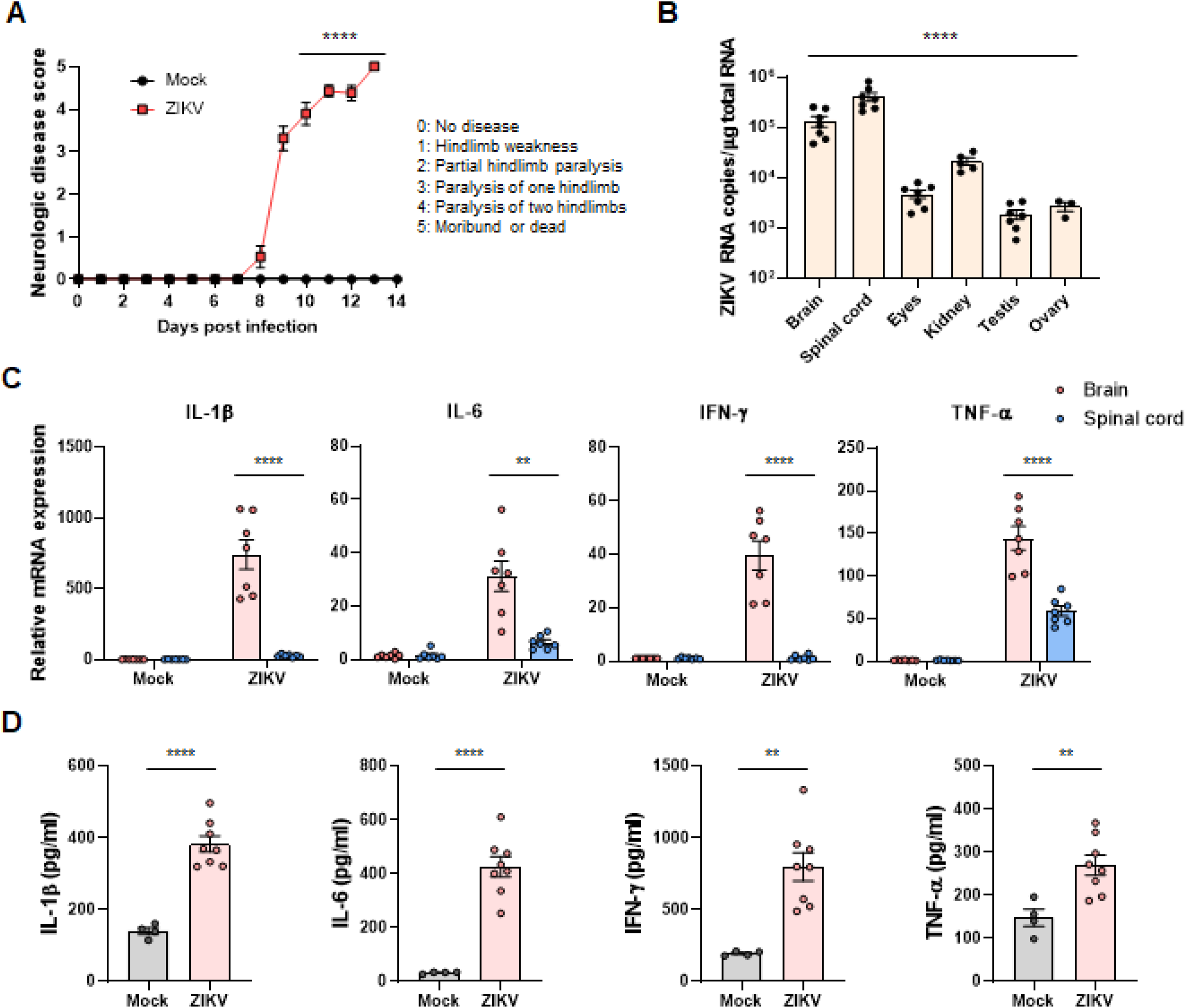
ZIKV infection of the brain elicits pro-inflammatory responses in *Ifnar1*^−/−^ mice. (A) *Ifnar1*^−/−^ mice were subcutaneously infected with 10^3^ PFU of ZIKV or PBS (mock). They were monitored daily for neurological disease score as described in Materials and Methods. (B) Viral RNA levels in the brain, spinal cord, eyes, kidney, testes, and ovary were assessed using RT-qPCR at 6 dpi. (C) RNA extracted from the brain and spinal cord homogenates was used to assess mRNA levels of proinflammatory cytokines, including IL-1β, IL-6, IFN-γ, and TNF-α, by RT-qPCR at 6 dpi. (D) The brain homogenates were used to measure protein levels of proinflammatory cytokines by ELSIA at 6 dpi. Statistically significant differences between the groups were determined using multiple two-tailed t-tests (A), one-way analysis of variance (ANOVA; B), and Student’s t-test (C, D); \**P* < 0.05; \*\**P* < 0.01; \*\*\**P* < 0.001; \*\*\*\**P* < 0.0001. Bars indicate mean ± SEM.

### Distinct transcriptional signatures and gene expression changes in the brain of ZIKV-infected *Ifnar1*^−/−^ mice

To assess the effects of ZIKV infection on gene expression in the mouse brain, we performed RNA-seq on brain homogenates of ZIKV-infected *Ifnar1*^−/−^ mice at 0, 3, and 6 dpi. Genes with adjusted *P*-value < 0.05 were considered differentially expressed genes (DEGs). We identified 930 DEGs at 6 dpi (Up-regulated: 546; Down-regulated: 384) that were differentially expressed compared to those at 0 dpi whereas there were 56 DEGs at 3 dpi (Up-regulated: 19; Down-regulated: 37). The volcano plot for DEGs at 6 dpi versus 0 dpi showed that several highly significant DEGs associated with IFN signaling were upregulated (Fig. 2A). According to the enrichment analysis of the biological category of gene ontology (GO), DEGs at 6 dpi were highly enriched in the cytokine-mediated signaling pathway, inflammatory responses, neutrophil-mediated immunity (Fig. 2B). The upregulation of DEGs involved in the immune response was most conspicuous at 6 dpi (Fig. 2C). Particularly, genes of proinflammatory cytokines (Il1b, Il6, and Tnf) and complement components (C1qa, C3, and C4b) in the inflammatory response were upregulated at 6 dpi (Fig. 2D). In addition, genes in the IFN signaling pathway (Oas1a, Oas2, Stat1, Stat2, Irf1, and Irf7) were upregulated at 6 dpi (Fig. 2E). Collectively, RNA-seq data from the brains of ZIKV-infected mice showed distinct immune and inflammatory signatures at 6 dpi.

**Fig. 2.**
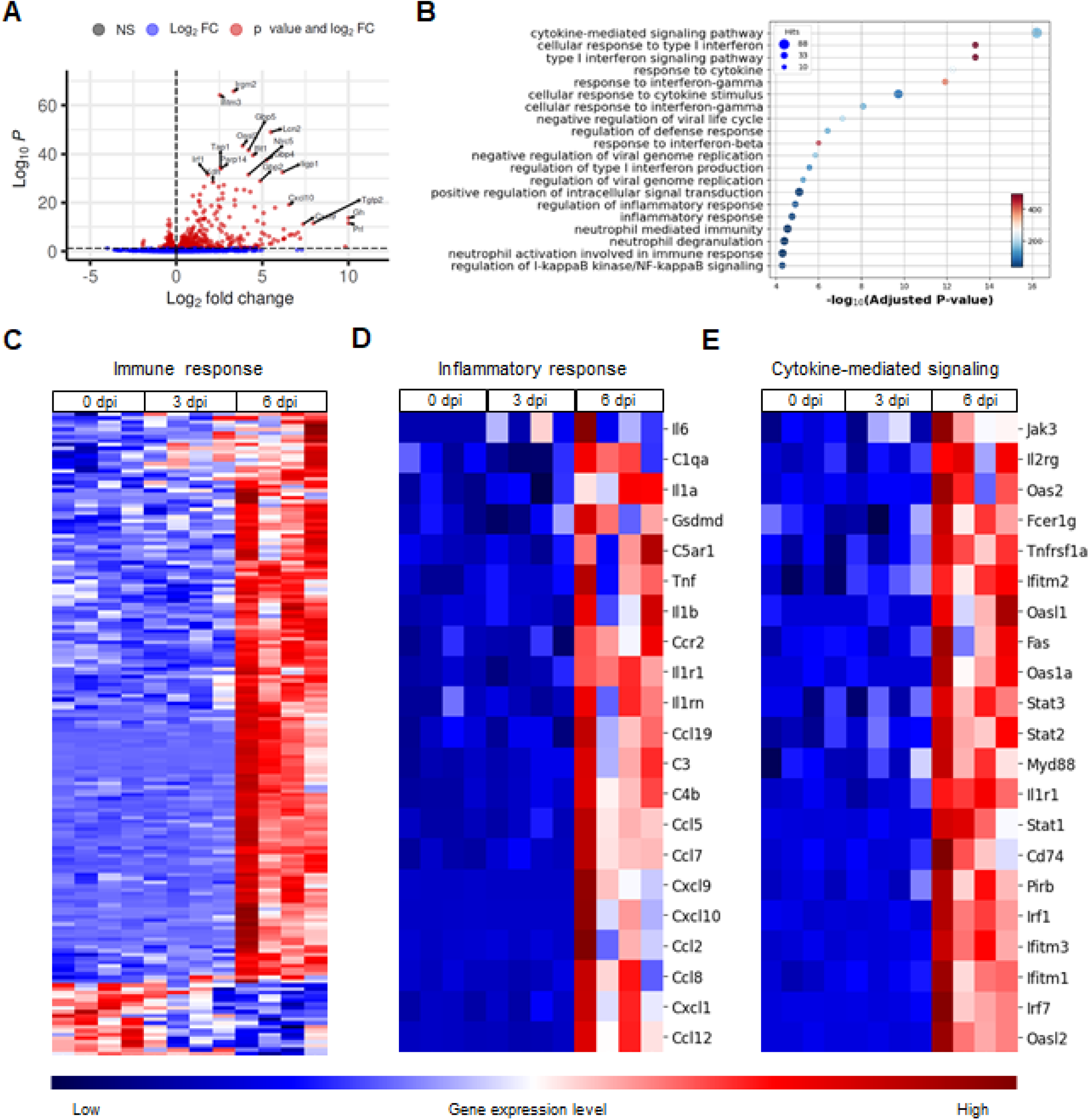
RNA-seq analysis of the brain in ZIKV-infected mice. *Ifnar1*^−/−^ mice were infected with 10^3^ PFU of ZIKV (n = 4 per indicated dpi). RNA extracted from the brain homogenates was used for RNA-seq analysis. (A) A volcano plot comparing DEGs from samples taken at 6 dpi versus 0 dpi. (B) GO enrichment analysis of biological process terms enriched in upregulated genes from comparisons of mice at 6 dpi versus 0 dpi. Terms were ranked by adjusted *P* value. (C–E) Heat maps of DEGs during ZIKV infection enriched in (C) immune response, (D) inflammatory response, and (E) cytokine-mediated signaling, identified through GO analysis. Gene expression levels in the heat maps are z score–normalized values determined from log_2_[CPM] values. NS: not significant; FC: fold change;

### ZIKV infection results in infiltration and proinflammatory activation of macrophages but not microglia in the *Ifnar1*^−/−^ mouse brain

To identify the immune cells in the brain that are responsible for the proinflammatory responses to ZIKV infection, we isolated the mouse brain after ZIKV infection. The brain homogenates were used to isolate immune cells in the brain, including microglia, macrophages, and lymphocytes, by 30 % and 70 % Percoll gradient centrifugation, followed by flow cytometry analysis of cell surface markers such as CD11b and CD45, as illustrated in Fig. 3A. The isolate consisted of three populations, namely lymphocytes (CD11b^-^and CD45^High^), macrophages (CD11b^+^ and CD45^High^), and microglia (CD11b^+^ and CD45^Low^), and their population was altered by ZIKV infection (Fig. 3B). While the number of microglia or lymphocytes in the brain did not change, that of macrophages increased approximately four-fold in response to ZIKV infection, possibly indicating macrophage brain infiltration (Fig. 3C–E). This pattern of alteration by ZIKV infection was different from that obtained by SARS-CoV-2 infection in our previous study, in which microglia were significantly depopulated (36). Next, we analyzed the IL-1β, IL-6, and TNF-α responses in each of the three populations (Fig. 4A). Interestingly, only the numbers of IL-1β-positive macrophages and lymphocytes, but not those of microglia, dramatically increased (Fig. 4B–F). These results suggest a role for macrophages and lymphocytes in IL-1β-mediated inflammation in the brains of ZIKV-infected mice.

**Fig. 3.**
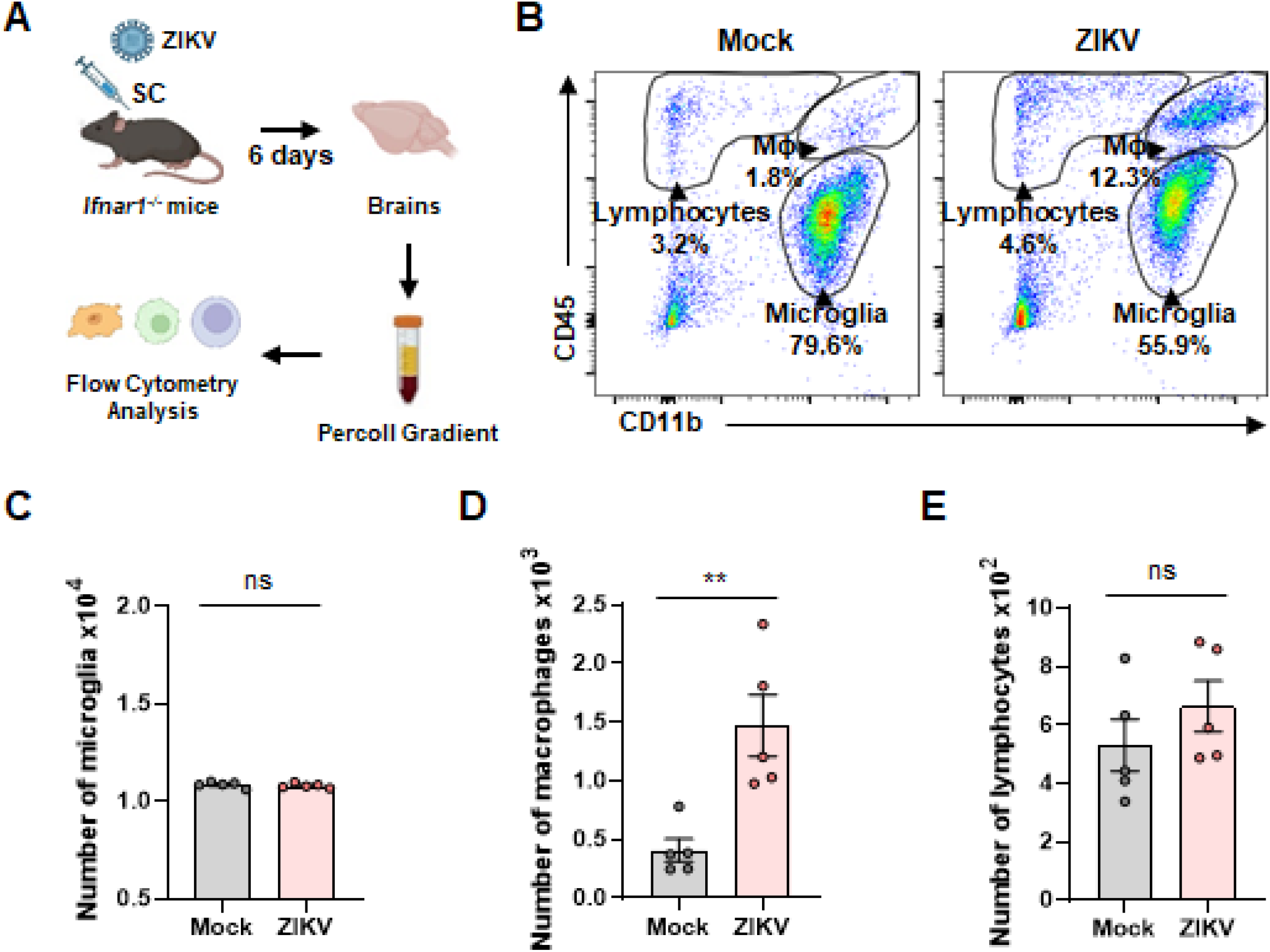
Macrophage infiltration into the brain of ZIKV-infected mice. (A) Schematic of the experiment for B to E, created with BioRender.com. *Ifnar1*^−/−^ mice were infected with 10^3^ PFU of ZIKV (n = 4) or PBS (mock; n = 4). At 6 dpi, brains of mock or ZIKV-infected mice were extracted and used for Percoll gradient centrifugation to isolate mononuclear cells for the flow cytometry analysis. The cellular surface of isolated mononuclear cells was stained with CD11b and CD45 antibodies. (B) Representative flow plot gated on leukocytes shows gating for microglia (MI, CD11b^+^, CD45^Low^), macrophages (Mϕ, CD11b^+^, CD45^High^), and lymphocytes (Lym, CD11b^-^, CD45^High^). (C–E) Bar graphs show the number of microglia (C), macrophages (D), and lymphocytes (E) isolated per brain at 6 dpi. Statistically significant differences between the groups were determined using Student’s t-test (C–E); ns, not significant; \*\**P* < 0.01; Bars indicate mean ± SEM.

**Fig. 4.**
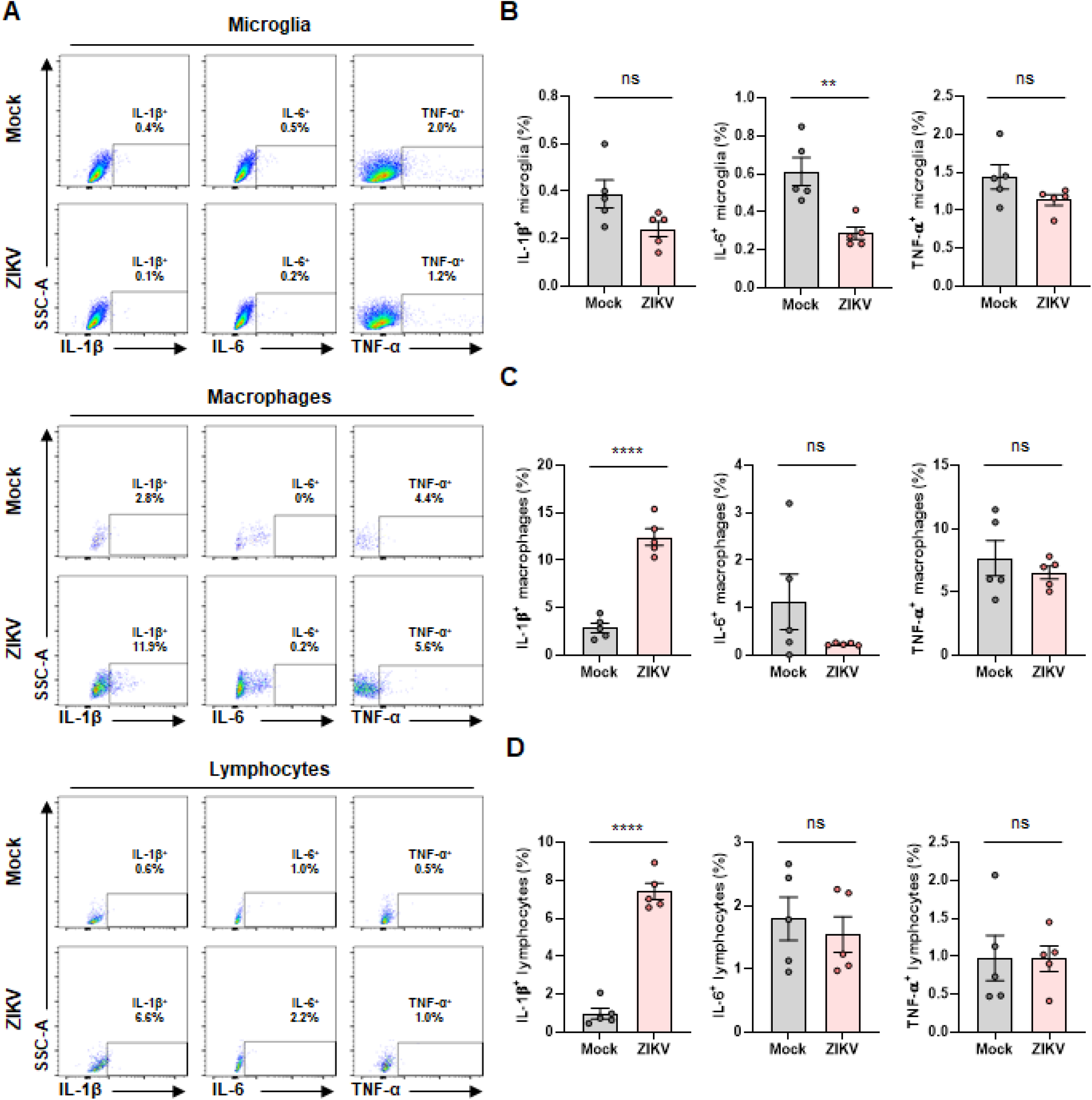
Proinflammatory activation of macrophages and lymphocytes by ZIKV infection in the mouse brain. (A) Representative flow plots gated on microglia (upper), macrophages (middle), and lymphocytes (lower) to show expression levels of proinflammatory cytokines, including IL-1β, IL-6, and TNF-α. (B–D) Bar graphs indicate the percentage of activated microglia (B), macrophages (C), and lymphocytes (D) expressing IL-1β, IL-6, and TNF-α. Statistically significant differences between the groups were determined using Student’s t-test (B–D); ns, not significant; \*\**P* < 0.01; \*\*\*\**P* < 0.0001. Bars indicate mean ± SEM.

We then investigated whether IL-1β expression in macrophages and lymphocytes was mediated by direct ZIKV infection. To address this, we used an anti-flavivirus envelope protein antibody (4G2) to stain ZIKV-infected cells along with IL-1β in the three isolated populations (Fig. 5A). As previous studies have demonstrated that microglia and macrophages are susceptible to ZIKV infection (18, 37-39), we observed that they were infected with ZIKV whereas lymphocytes were not infected (Fig. 5B–D). While ZIKV infection of microglia in *Ifnar1*^−/−^ mice did not lead to microglial activation (Fig. 4B, 5E), infection of macrophages resulted in the induction of IL-1β (Fig. 4C, 5F). Lymphocytes showed elevated IL-1β levels, but they were not infected with ZIKV (Fig. 4D, 5G). These findings demonstrate that IL-1β induction in the brains of ZIKV-infected *Ifnar1*^−/−^ mice was likely due to ZIKV-infected macrophages and lymphocytes, but not microglia.

**Fig. 5.**
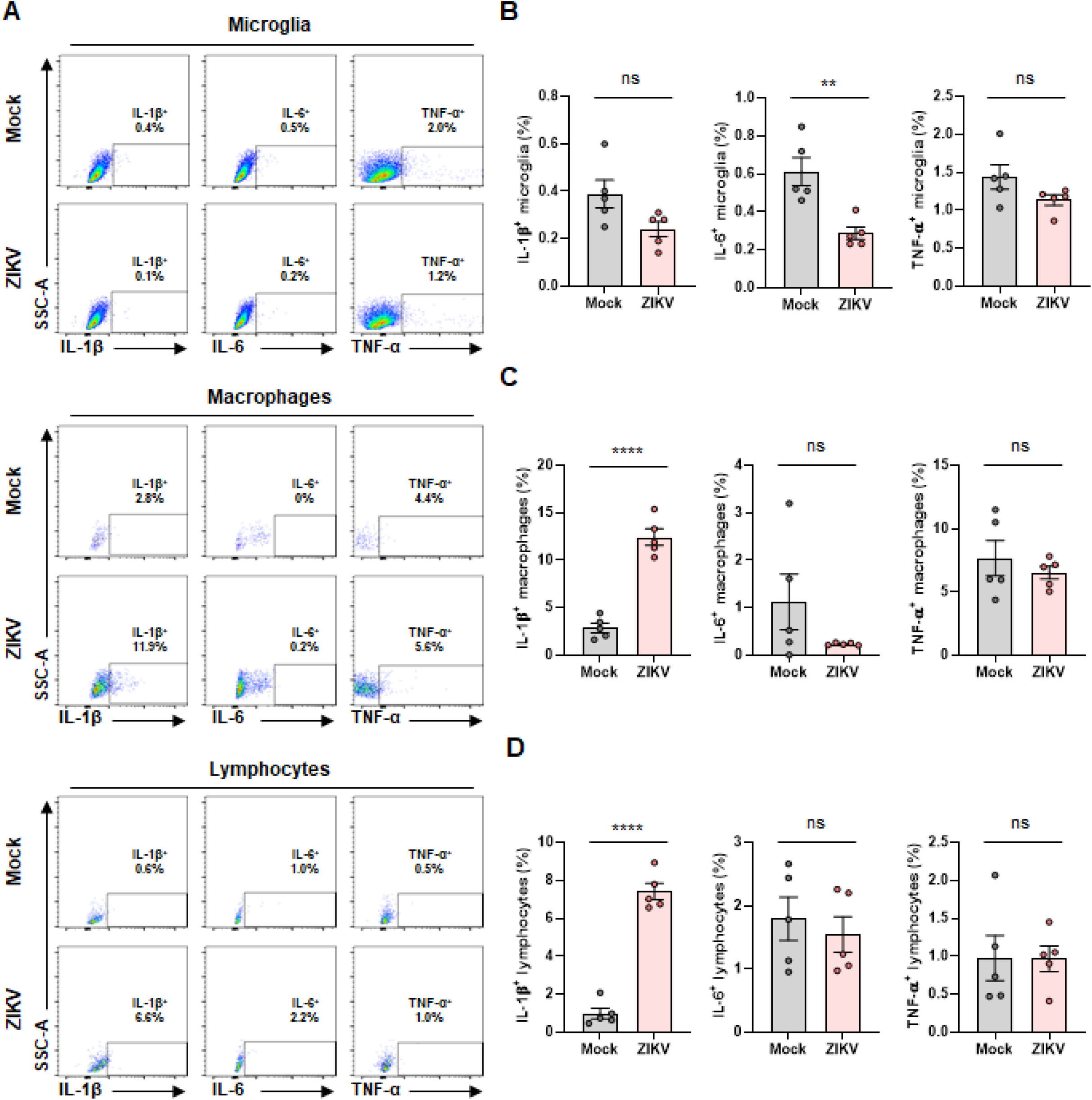
Microglia and macrophage infection by ZIKV in the mouse brain. *Ifnar1*^−/−^ mice were infected with 10^3^ PFU of ZIKV (n = 4) or PBS (Mock; n = 4). At 6 dpi, isolated mononuclear cells in brains of mock or ZIKV-infected mice were used for the flow cytometry analysis. (A) Representative flow plots gated on microglia (upper), macrophages (middle), and lymphocytes (lower) to show expression levels of ZIKV envelop protein and IL-1β. (B–D) Bar graphs indicate median fluorescence intensities of ZIKV envelop protein-positive microglia (B), macrophages (C), and lymphocytes (D). (E–G) Bar graphs indicate the percentage of activated microglia (E), macrophages (F), and lymphocytes (G) expressing IL-1β. Statistically significant differences between the groups were determined using Student’s t-test (B–G); ns, not significant; \**P* < 0.05; \*\*\*\**P* < 0.0001. Bars indicate mean ± SEM.

### ZIKV infection of THP-1 cells induces IL-1β secretion and inflammatory cell death

Next, we used human monocyte THP-1 cells to confirm our *in vivo* observations. Given the key role of NLRP3 inflammasome in innate immune responses by activating caspase-1 to promote IL-1β secretion and pyroptosis (40), we examined whether ZIKV infection stimulates IL-1β secretion through NLRP3 inflammasome activation in THP-1 cells. When THP-1 cells were infected with ZIKV at a multiplicity of infection (MOI) of 0.1, the viral RNA and envelope protein levels were increased in a time-dependent manner, as shown by RT-qPCR and western blotting, respectively (Fig. 6A, 6B). ELISA revealed that, compared to the mock infection, ZIKV infection significantly augmented IL-1β secretion over time (Fig. 6C). In addition to the increase in IL-1β secretion, cell death was promoted by ZIKV infection in THP-1 cells, possibly due to pyroptosis (Fig. 6D). Consequently, we determined whether caspase-1 and GSDMD cleavage resulting from activation of the NLRP3 inflammasome were induced by ZIKV infection. Caspase-1 and Gasdermin D (GSDMD) cleavage was observed from 1 dpi, followed by IL-1β maturation and secretion (Fig. 6E). Thus, ZIKV infection of THP-1 cells activates the NLRP3 inflammasome and consequently induces IL-1β maturation and secretion.

**Fig. 6.**
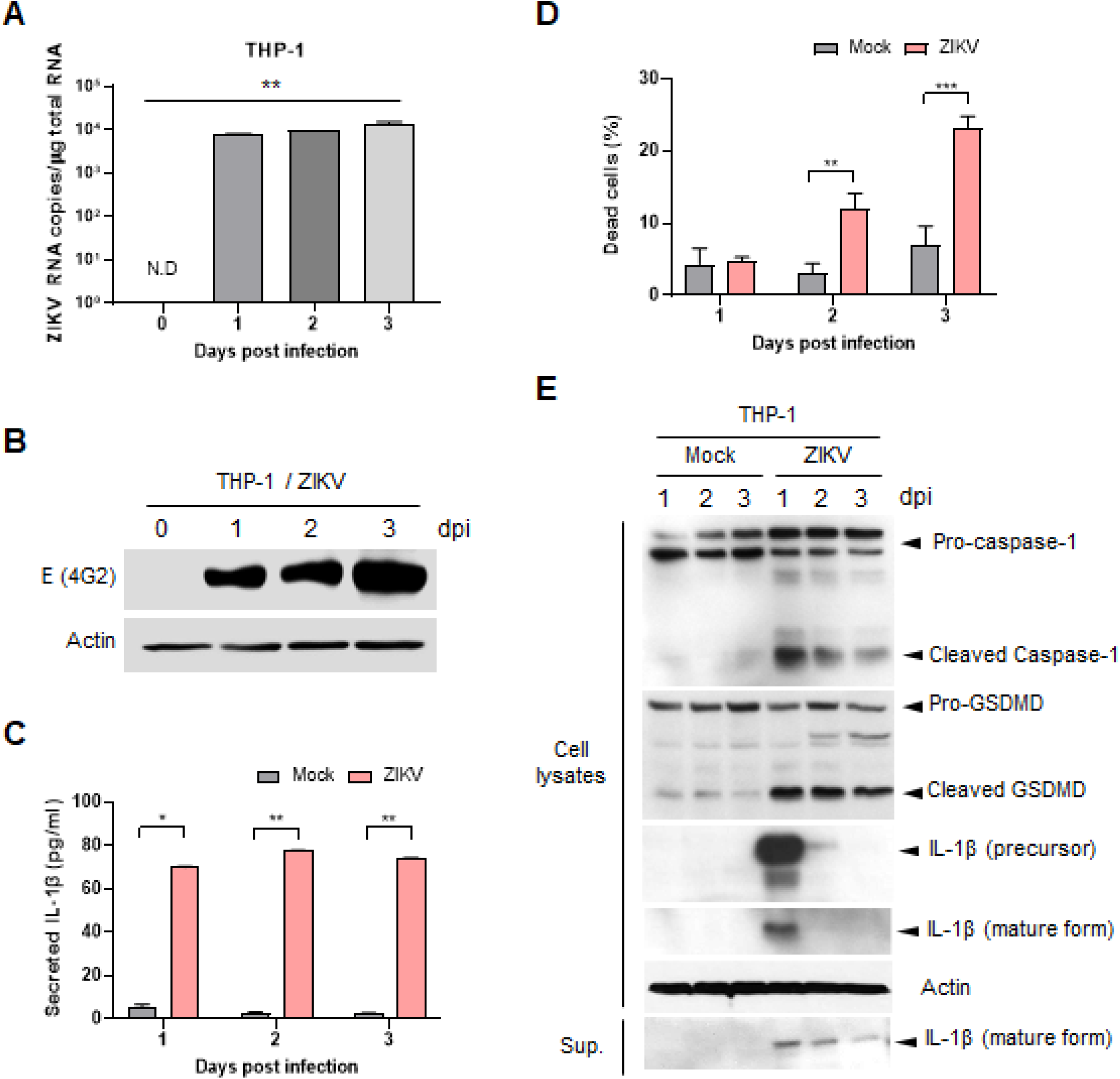
ZIKV infection induces IL-1β secretion and inflammatory cell death in THP-1 cells. THP-1 cells were infected with ZIKV at 0.1 MOI. (A) Viral loads in ZIKV-infected THP-1 cells were determined by RT-qPCR at 0, 1, 2, and 3 dpi. (B) ZIKV envelop protein in cell lysates were determined by western blot. Actin served as the loading control. (C) IL-1β levels in the cell culture media were measured by ELISA at 1, 2, and 3 dpi. (D) Dead cells were analyzed using Trypan Blue dye exclusion at 1, 2, and 3 dpi. (E) Pro-caspase-1, cleaved caspase-1, GSDMD, cleaved GSDMD, pro-IL-1β, and mature IL-1β in cell lysates or in supernatants were determined by western blot. Actin served as the loading control. Statistically significant differences between the groups were determined using one-way analysis of variance (ANOVA; A) and Student’s t-test (C and D); \**P* < 0.05; \*\**P* < 0.01; \*\*\**P* < 0.001. Bars indicate mean ± SEM.

### The increase in C3 levels by ZIKV infection is mediated by IL-1β signaling

Previous studies have revealed that IL-1β and other proinflammatory cytokines induce the transcription factor CCAAT/enhancer binding protein β (C/EBP-β); this transcription factor directly activates the promoter of C3, which plays a crucial role in the activation of the complement system and contributes to innate immune responses (27, 41, 42). To validate the role of C/EBP-β in IL-1β-mediated induction of C3 expression by ZIKV infection in THP-1 cells, we infected cells with ZIKV at 5 MOI and analyzed the activation of P38, Erk 1/2, and C/EBP-β using western blotting. Indeed, ZIKV infection induced C/EBP-β expression and its activation through phosphorylation, which was conducted by activated p38 but not Erk 1/2, resulting in the elevation of C3 expression (Fig. 7A). C3 induction was suppressed by C/EBP-β knockout in ZIKV-infected THP-1 cells (Fig. 7B). Diacerein, an inhibitor of IL-1β production (43), and an IL-1R antagonist effectively reduced the ZIKV-mediated C3 induction in a dose-dependent manner, suggesting the involvement of IL-1β in the induction of C3 gene expression by ZIKV infection (Fig. 7C, 7D). The reduction of C3 secretion by Diacerein and the IL-1R antagonist was confirmed by ELISA (Fig. 7E, 7F). Therefore, we demonstrated that ZIKV infection in THP-1 cells promotes C/EBP-β expression through IL-1β induction, eventually leading to increased C3 levels.

**Fig. 7.**
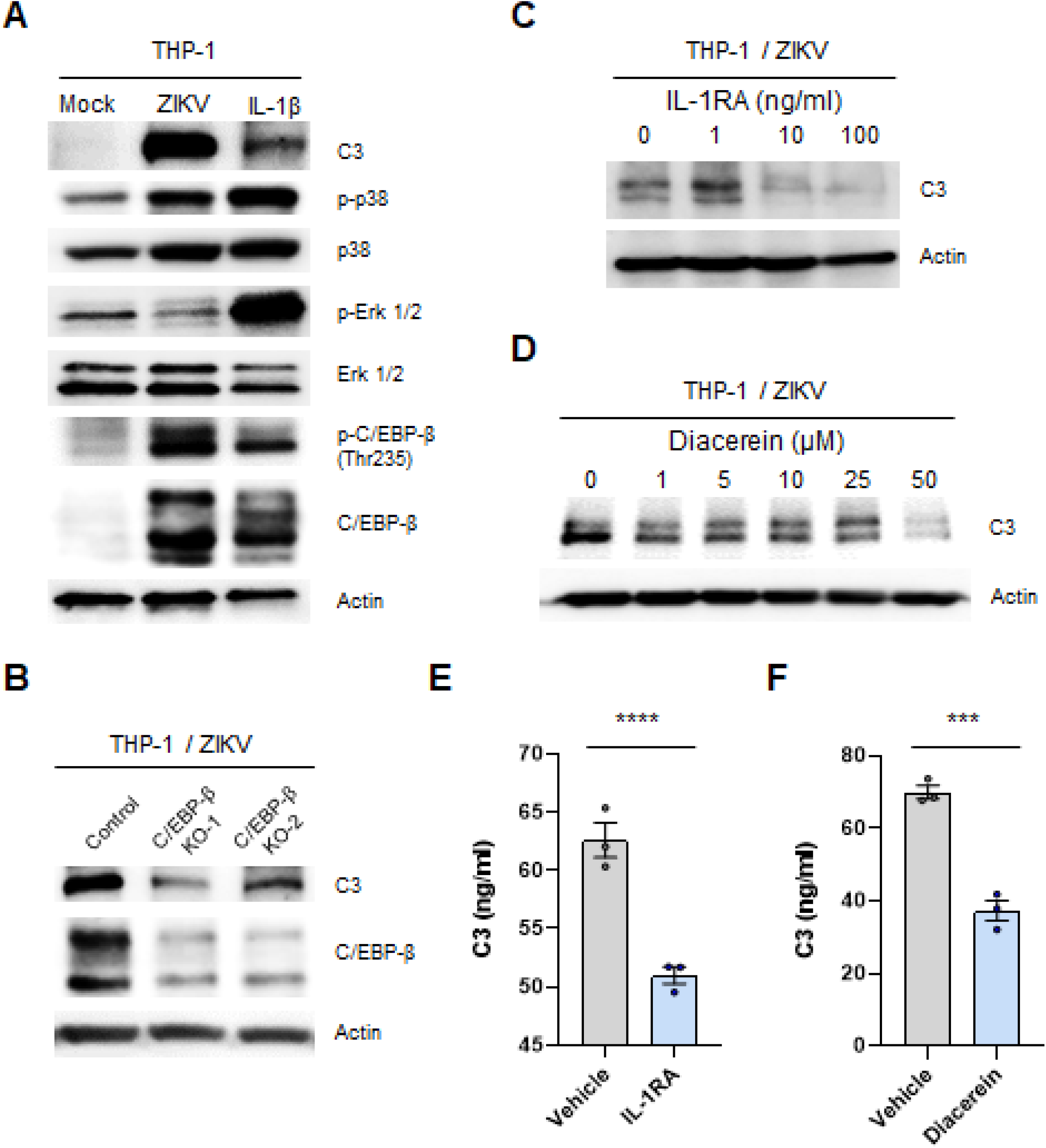
ZIKV infection induces C3 expression through IL-1β signaling in THP-1 cells. (A) THP-1 cells were infected with ZIKV at 5 MOI or treated with 10 ng/ml IL-1β. At 2 dpi, C3, phospho-p38, p38, phospho-Erk 1/2, Erk 1/2, phospho-C/EBP-β (Thr235), and C/EBP-β levels in cell lysates were assessed by western blot. Actin served as the loading control. (B) C/EBP-β knockout THP-1 cell lines (C/EBP-β KO-1 and C/EBP-β KO-2) were infected with ZIKV at 5 MOI. C3 and C/EBP-β proteins in cell lysates were quantitatively analyzed by western blot. Actin served as the loading control. (C–F) THP-1 cells were infected with ZIKV at 5 MOI, followed by treatment with an IL-1R antagonist as indicated (C) or 100 ng/ml (E), or with Diacerein as indicated (D) or 50 μM (F). C3 levels in cell lysates were assessed by western blot (C and D). Actin served as the loading control. Secreted C3 levels in supernatants were measured by ELISA (E and F). Statistically significant differences between the groups were determined using Student’s t-test (E and F); \*\*\**P* < 0.001; \*\*\*\**P* < 0.0001. Bars indicate mean ± SEM.

In the brains of *Ifnar1*^−/−^ mice, the induction and activation of C/EBP-β by ZIKV infection were determined at 6 dpi using RT-qPCR and western blot analysis (Fig. 8A, 8B). The increased expression and activation of C/EBP-β led to the induction of C3 mRNA and protein expression as determined by RT-qPCR and ELISA, respectively (Fig. 8C, 8D). We also detected C3 induction in mouse sera following ZIKV infection, which indicated systemic C3 induction. To evaluate the functional consequences of C3 induction and complement activation by ZIKV infection, we determined the C5a levels in the brain. As expected, the C5a levels in the brains of ZIKV-infected mice were significantly increased at 6 dpi (Fig. 8F). Taken together, our findings indicate that the induction of C3 and complement activation by ZIKV infection is mediated by the IL-1β signaling pathway.

**Fig. 8.**
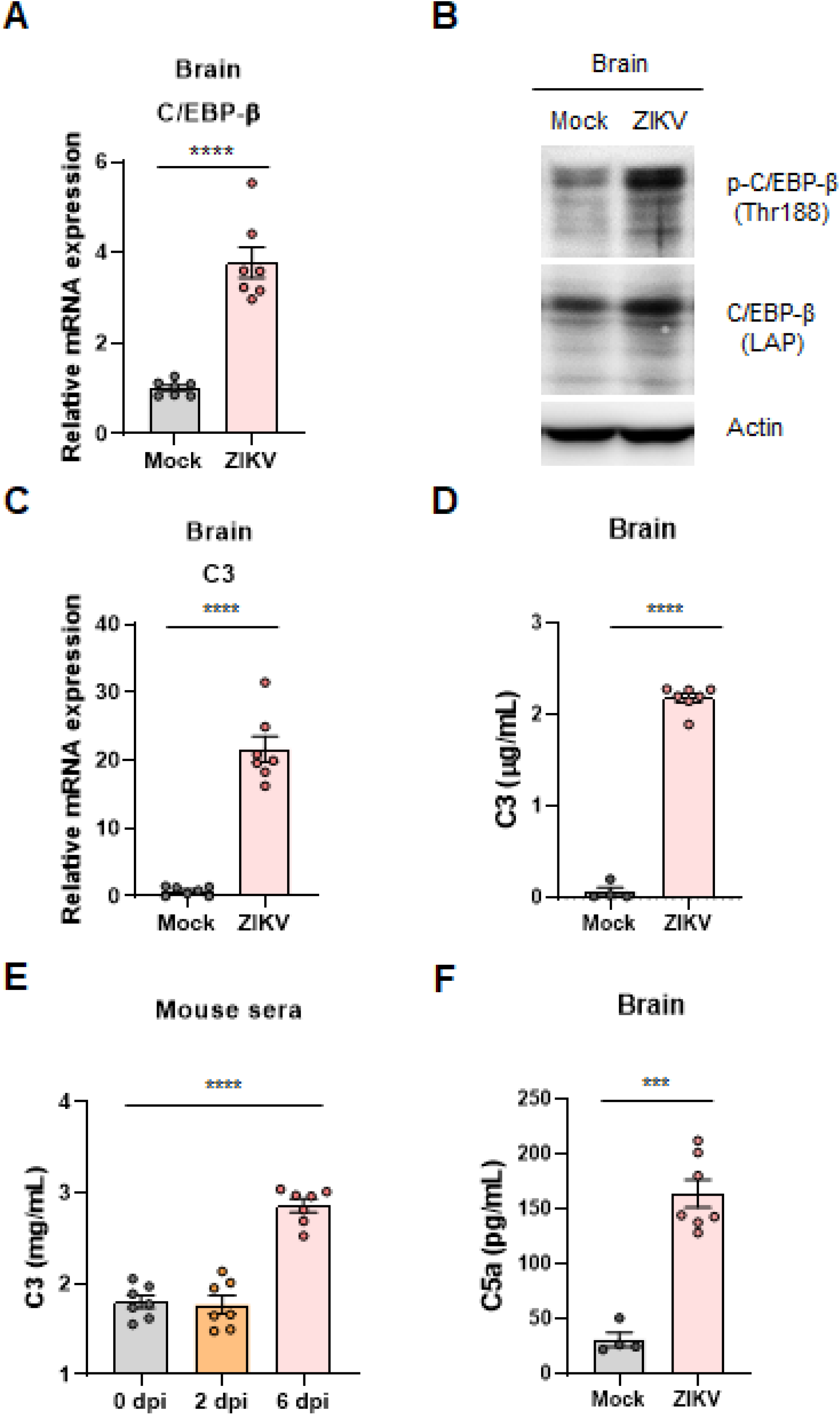
ZIKV infection promotes C/EBP-β and C3 expression and activation in the *Ifnar1*^−/−^ mice. *Ifnar1*^−/−^ mice were infected with 10^3^ PFU of ZIKV (n = 7) or PBS (mock; n = 4 to 7). (A) C/EBP-β mRNA levels in the brain homogenates were determined by RT-qPCR. (B) Phospho-C/EBP-β (Thr188) and C/EBP-β (LAP) protein levels in lysates of the brain homogenates were assessed by western blot. Actin served as the loading control. (C) C3 mRNA levels in the brain homogenates were determined by RT-qPCR. (D) C3 protein levels in lysates of the brain homogenates were assessed by ELISA. (E) C3 protein levels in mouse sera were measured by ELISA. (F) C5a protein levels in lysates of the brain homogenates were quantitatively analyzed by ELISA. Statistically significant differences between the groups were determined using Student’s t-test (A, C, D, and F) and one-way analysis of variance (ANOVA; E); \*\*\**P* < 0.001; \*\*\*\**P* < 0.0001. Bars indicate mean ± SEM.

## Discussion

To date, the neuropathogenesis of ZIKV infection remains unclear. Understanding the neuropathogenesis of ZIKV infection would help to understand ZIKV-associated neurological diseases. ZIKV infects human fetal brain microglia and macrophages (19). This induces high levels of proinflammatory cytokines, which are implicated in ZIKV-derived neuroinflammation. C3 is induced in most neurodegenerative diseases and is upregulated by proinflammatory cytokines such as IL-1β (24-27). Furthermore, IL-1β and C3 expression is increased in different viral infections of the CNS (28, 44). However, whether IL-1β and C3 expression in the brain is affected by ZIKV infection remains to be investigated. Here, we used adult *Ifnar1*^−/−^ mice as a ZIKV infection mouse model to examine proinflammatory responses in the CNS upon ZIKV infection. We demonstrated that ZIKV infection of the brain in this animal model augments IL-1β expression in infiltrating macrophages and elicits IL-1β-mediated inflammation, leading to the destructive consequences of neuroinflammation.

Neurons are targets in the CNS for ZIKV, JEV, and WNV infection (45-47). They are likely the first responders to immune responses against neurotropic viral infections, producing type I IFNs and expressing MHC class I molecules (48, 49). Upon initiation of an inflammatory response in the CNS, proinflammatory cytokines are critical for recruitment of monocytes to the CNS (44). When we infected the *Ifnar1*^−/−^ mice with ZIKV, increased pro-inflammatory cytokine levels and macrophage infiltration were observed in the brain at 6 dpi (Fig. 1C, 1D, 3B, 3D). These results imply that ZIKV infection can elicit neuronal inflammation and, consequently, monocyte infiltration in this animal model.

Microglia and macrophages have been found to be of the same origin (50) and have similar functions, such as production of inflammatory mediators (51). However, they play different roles in the brain. Microglia may protect the injured brain, whereas macrophages concurrently damage the brain (52). In this study, although both microglia and macrophages were infected with ZIKV at 6 dpi (Fig. 5B, 5E), macrophages showed proinflammatory activation with upregulated IL-1β expression, but microglia were not observed in the brains of infected *Ifnar1*^−/−^ mice (Fig. 4B, 4C, 5E, 5F). Proinflammatory activated macrophages can accelerate the number of circulating immune cells and increase their infiltration into the brain, also playing critical roles in pathophysiological processes. Further studies are warranted to determine why ZIKV-infected microglia were not activated in the brains of *Ifnar1*^−/−^ mice.

In ZIKV-GBS, particularly the acute inflammatory demyelinating polyneuropathy variant, cytokine-mediated inflammation and macrophage activation may lead to peripheral nerve injury, and complement activation may be associated with demyelinating neuropathies (8, 53). Given that IL-1 signaling is considered the upper hierarchical cytokine signaling cascade in the CNS (29), ZIKV infection of the brain in *Ifnar1*^−/−^ mice induces IL-1β expression in infiltrating macrophages (Fig. 3D, 4C, 5F) and elicits IL-1β-mediated inflammation and macrophage activation, which can be associated with the destructive consequences of neuroinflammation, such as peripheral neuropathy. Our findings also suggest that ZIKV-induced IL-1β secretion might lead to the expansion of encephalitogenic T cells (54) and pyroptosis in CNS peripheral myeloid and lymphoid cells; these cells can mediate neuroinflammation in multiple CNS diseases (55).

Complement is thought to have a protective effect, but exaggerated or insufficient activation of the complement system can cause neuropathies and contribute to neurodegeneration and neuroinflammation (24, 56). ZIKV infection upregulated C3 expression through IL-1β-mediated signaling (Fig. 7 and 8), possibly disrupting the balance of the complement system, which can mediate myelin phagocytosis by macrophages (56). The anaphylatoxins (C3a, C4a, and C5a) produced during complement activation play a major role in the pathogenesis of inflammatory disorders, including ischemia/reperfusion injury, and are involved in various neurodegenerative disorders (57). An increase in C5a in the cerebrospinal fluid was also detected during the exacerbation of neuromyelitis (58). In this study, we observed an increase in C5a levels in the brains of ZIKV-infected *Ifnar1*^−/−^ mice (Fig. 8F), suggesting its contribution to peripheral neuropathies (59).

In summary, ZIKV infection of the brain induced the expression of proinflammatory cytokines, including IL-1β, IL-6, IFN-γ, and TNF-α, in *Ifnar1*^−/−^ mice. RNA-seq analysis revealed that the expression of genes involved in immune responses to viral infection and inflammatory responses in the brains of these mice was significantly upregulated by ZIKV infection. Notably, infiltration and pro-inflammatory activation of macrophages, but not microglia, were observed at 6 dpi in the ZIKV-infected *Ifnar1*^−/−^ mice. ZIKV infection of THP-1 cells induced pyroptosis, increased IL-1β release, and consequently increased C3 expression. Increases in C3 and C5a levels in the brains of infected mice were confirmed. Overall, our data suggest that ZIKV infection of the brains of *Ifnar1*^−/−^ mice augments IL-1β expression in infiltrating macrophages and elicits IL-1β-mediated inflammation through macrophage activation, which can be associated with the destructive consequences of neuroinflammation.

## Materials and methods

### Cells, plasmids, and virus

THP-1 (TIB-202), a human leukemia monocytic cell line, and the ZIKV PRVABC59 strain (VR-1843; GenBank <u>KU501215</u>) were purchased from the American Type Culture Collection (ATCC; Manassas, VA, USA). This cell line was maintained in RPMI-1640 medium (Cytiva, Marlborough, MA, USA) containing 10 % fetal bovine serum (FBS) (Gibco, Waltham, MA, USA) and 1 % penicillin/streptomycin (Gibco). This virus was propagated in Vero cells (CCL-81; ATCC). Virus titers were determined using plaque assay as described previously (60).

For CRISPR/Cas9-mediated knockout of C/EBP-β, we constructed CRISPR/Cas9 plasmids using the LentiCRISPRv2-puro vector (Addgene #98290) (61, 62). LentiCRISPRv2-C/EBP-β plasmids were generated by inserting hybridized oligonucleotides (#1:5’-CTCTTCTCCGACGACTACGG-3’ and 5’-CCGTAGTCGTCGGAGGAGAG-3’; #2:5’-GGCCAACTTCTACTACGAGG-3’ and 5’-CCTCGTAGTAGAAGTTGGCC-3’) into the BsmBI restriction sites. Lentivirus was produced from 70–80 % confluent HEK293T cells transfected with 0.2 μg lentiviral plasmid, 0.5 μg pVSV-G (Addgene #8454), and 2.3 μg psPAX2 (Addgene #12260) using 6 μl of lipofectamine 2000 (Invitrogen). The lentiviral supernatant was harvested 48 h later and used for the transduction of THP-1 cells by spinoculation. After 48 h post transduction, the cells were selected with 1 μg/ml puromycin (Gibco) for a further seven days.

### Drug treatment and cell viability assay

Cells (1 × 10^6^ cells per well) were plated into six-well plates and inoculated with 0.1 or 5 MOI ZIKV in RPMI-1640 medium the next day. After 1 h incubation, cells were treated with Diacerein (D9302; Sigma-Aldrich, St. Louis, MO, USA) or IL-1RA (SRP3327, Sigma-Aldrich). At the indicated time points, the cells were washed with PBS by cell pelleting for cell lysates and RNA extraction. The cells were lysed in radioimmunoprecipitation (RIPA) buffer (Thermo Fisher Scientific, Waltham, MA, USA). Total cellular RNA was extracted using an RNeasy Mini Kit (QIAGEN, Hilden, Germany). To determine cell viability, cells were monitored using Trypan Blue dye exclusion.

### Mice

*Ifnar1*^−/−^ mice were purchased from B&K Universal Ltd. (Hull, UK). Mice were bred and housed in a BSL-2 animal facility at the Korea Research Institute of Chemical Technology (KRICT). Male and female mice were between six and eight weeks of age at the initiation of all the experiments. All protocols were approved by the Institutional Animal Care and Use Committee (Protocol ID 8A-M6, IACUC ID 2021-8A-02-01, and 2021-8A-03-03). Viral inoculations (10^3^ pfu) were performed subcutaneously under anesthesia using isoflurane in the BSL-2 animal facility, and all efforts were made to minimize animal suffering. Body weight was measured daily post-infection. Neurological symptoms were scored from 0 to 5 as follows: 0, no symptoms; 1, hindlimb weakness; 2, partial hindlimb paralysis; 3, paralysis of one hindlimb; 4, paralysis of both hindlimbs; and 5, moribund or dead.

### Mononuclear cell isolation

Mononuclear cells isolation in the mouse brain was performed following the protocols as previously described (36, 63, 64). Mock- or ZIKV-infected mice were anesthetized with isoflurane, followed by perfusion with 10 ml of cold 1× DPBS (Gibco) into the left ventricle to remove blood from the tissues. The brains were transferred to a six-well plate containing cold Hanks’ balanced salt solution (HBSS) (Gibco), and the plates were kept on ice. The generation of brain cell suspensions by a 70 μm pore sized-cell strainer (SPL, Gyeonggi-do, South Korea) was made in 10 ml per brain of digestion cocktail containing 0.5 mg/ml DNase I (Roche, Basel, Switzerland) and 1 mg/ml Collagenase A (Roche) in HBSS. The suspension was incubated at room temperature for 30 min, followed by centrifugation for 7 min at 300 × g and 18 °C. The cell pellet was resuspended in 30 % Percoll (Sigma-Aldrich) in HBSS and then slowly layered over 70 % Percoll in HBSS in a 15 ml conical tube. Approximately 2 ml of the interphase volume was collected into a new tube after gradient centrifugation for 40 min at 200 × g at 18 °C. Isolated mononuclear cells were washed three more times in a volume of 500 μL of HBSS containing 0.01 M HEPES (Gibco), using a microcentrifuge for 7 min at 600 × g at 4 °C.

### Flow cytometry analysis

Isolated brain mononuclear cells from the mock or ZIKV-infected mice in cell staining buffer (PBS with 1 % FBS and 0.09 % NaN_3_) were stained for 30 min with fluorescence-conjugated antibodies, namely, Brilliant Violet 421 anti-mouse/human CD11b Antibody (101236, BioLegend, San Diego, CA, USA), PE/Cyanine7 anti-mouse CD45 Antibody (103114, BioLegend), APC anti-mouse TNF-α Antibody (506307, BioLegend), FITC anti-mouse IL-6 Monoclonal Antibody (MP5-20F3) (11-7061-82, eBioscience, San Diego, CA, USA), PE anti-mouse IL-1β Antibody (12-7114-82, eBioscience), and Alexa Fluor 488 anti-flavivirus group antigen antibody (4G2) (NBP2-52709AF488, Novus Biologicals, Englewood, CO, USA). The cells were then analyzed using a FACSAria III sorter (BD Biosciences, San Jose, CA, USA), and data were analyzed using FlowJo software (BD Biosciences). All fluorochromes were compensated for.

### RT-qPCR

Quantitative RT-PCR (QuantStudio 3, Applied Biosystems, Foster City, CA, USA) was performed using one-step Prime script III RT-qPCR mix (Takara Bio, Shiga, Japan). The viral RNA of ZIKV NS3 was detected by customized probe-based qPCR assay (Integrated DNA Technologies, Coralville, IA, USA). The Il1b, Il6, Tnf, Ifng, Cebpb, and C3 genes were also detected using individual customized probes (Integrated DNA Technologies). The sequences of the qPCR probes and primers used in this study are listed in Table S1.

### RNA-seq and analysis

The sequencing library was prepared using the TruSeq Stranded mRNA Sample Prep Kit and sequenced on NovaSeq 6000 (Illumina, San Diego, CA, USA), yielding more than 6G bases of sequence for each sample. Adaptor sequences were removed from the sequenced reads using Cutadapt (version 3.1) (65) and aligned to the hybrid reference genomes of humans (GRCh38.p13_ENS100) and ZIKV with the STAR aligner (version 2.7.6a) (66). Aligned reads were quantified at the gene level by HTSeq (version 0.13.5) (67) with “intersection-nonempty” mode. Genes with lower than five counts for the total count per gene were removed for further analysis. Differentially expressed genes were analyzed with DESeq2 (version 1.30.1) (68) using abs (log2 fold change) > 1 and adjusted P-value (Benjamini-Hochberg) < 0.01 as the cut-off. Multidimensional scaling analysis was performed with the clustermap function in the Python seaborn package (version 0.11.1) using genes with a mean FPKM > 1 among the samples and transformed to log2(FPKM+1). Over-representation analysis of the DEGs enriched in GO Biological Process 2018 was performed with EnrichR (69) and an adjusted P-value (Benjamini-Hochberg) < 0.05.

### ELISA

Culture supernatants collected from infected cells, brain homogenates, and mouse sera were used for the detection of IL-1β, IL-6, TNF-α, IFN-γ, C3, and C5a. The concentration of each of them was determined using an ELISA kit ([human IL-1β, K0331800; mouse IL-1β, K0331231; IL-6, K0331230; TNF-α, K0331186; IFN-γ, K0331138; Komabiotech, Seoul, Korea]; [mouse C3, ab263884; C5a, ab193718; Abcam, Cambridge, UK]), according to the manufacturer’s instructions.

### Western blotting

Proteins in the lysate were separated on a denaturing polyacrylamide gel and transferred to a polyvinylidene fluoride (PVDF) membrane (Merck Millipore, Burlington, MA, USA). The membrane was incubated with 5 % skim milk (BD Biosciences) in Tris-buffered saline with 0.1 % Tween 20 (TBST) buffer and the primary antibodies, namely, anti-IL-1β (GTX130021, GeneTex, Irvine, CA, USA), anti-GSDMDC1 (sc-81868, Santa Cruz Biotechnology), anti-β-actin (sc-47778, Santa Cruz Biotechnology), anti-Caspase-1 (3866S, Cell Signaling Technology), anti-pan-flavivirus E (4G2, purified in the lab), anti-C3 (ab200999, Abcam), anti-C/EBP-β (90081S, Cell Signaling Technology), anti-C/EBP-β (LAP) (3087S, Cell Signaling Technology), anti-phospho-C/EBP-β (Thr235) (3084S, Cell Signaling Technology), anti-p38 MAPK (9212S, Cell Signaling Technology), anti-phospho-p38 MAPK (9211S, Cell Signaling Technology), anti-p44/42 MAPK (9102S, Cell Signaling Technology), anti-phospho-p44/42 MAPK (4370S, Cell Signaling Technology). Horseradish peroxidase (HRP)-conjugated secondary antibodies from Bio-Rad and enhanced chemiluminescence (ECL) reagents (Thermo Fisher Scientific) were used for protein detection.

### Statistical analysis

All experiments were performed at least thrice. All data were analyzed using the GraphPad Prism 8.0 software (GraphPad Software, San Diego, CA, USA). P < 0.05 was considered statistically significant. Specific analysis methods are described in the figure legends.

## Acknowledgments

This work was supported by the National Research Foundation of Korea (NRF) grant funded by the Ministry of Education, Science, and Technology (MIST) of the Korean government (2020R1C1C1003379) and the National Research Council of Science & Technology (NST) grant funded by the Korean government (MSIP) (CRC-16-01-KRICT). The authors declare that they have no conflicts of interest.

## Conceptualization

G.U.J., S.L., and Y.-C.K.; methodology: G.U.J., S.L., and Y.-C.K.; investigation: G.U.J., S.L., D.Y.K., J.K., G.Y.Y., and J.K.; writing: G.U.J. and Y.-C.K..; review and editing: G.U.J., S.L., and Y-C.K.; funding acquisition: Y.-C.K.; and supervision: Y.-C.K.

The authors declare that they have no conflicts of interest.

## References

1. Ikejezie J, Shapiro CN, Kim J, Chiu M, Almiron M, Ugarte C, et al. Zika virus transmission—region of the Americas, May 15, 2015–December 15, 2016. Morbidity and Mortality Weekly Report. 2017;66(12):329.

2. de Araújo TVB, Rodrigues LC, de Alencar Ximenes RA, de Barros Miranda-Filho D, Montarroyos UR, de Melo APL, et al. Association between Zika virus infection and microcephaly in Brazil, January to May, 2016: preliminary report of a case-control study. The lancet infectious diseases. 2016;16(12):1356–63.

3. Mattar S, Ojeda C, Arboleda J, Arrieta G, Bosch I, Botia I, et al. Case report: microcephaly associated with Zika virus infection, Colombia. BMC infectious diseases. 2017;17(1):1–4.

4. Cao-Lormeau V-M, Blake A, Mons S, Lastère S, Roche C, Vanhomwegen J, et al. Guillain-Barré Syndrome outbreak associated with Zika virus infection in French Polynesia: a case-control study. The Lancet. 2016;387(10027):1531–9.

5. Mécharles S, Herrmann C, Poullain P, Tran T-H, Deschamps N, Mathon G, et al. Acute myelitis due to Zika virus infection. The Lancet. 2016;387(10026):1481.

6. Carteaux G, Maquart M, Bedet A, Contou D, Brugières P, Fourati S, et al. Zika virus associated with meningoencephalitis. New England Journal of Medicine. 2016;374(16):1595–6.

7. Soares CN, Brasil P, Carrera RM, Sequeira P, De Filippis AB, Borges VA, et al. Fatal encephalitis associated with Zika virus infection in an adult. Journal of Clinical Virology. 2016;83:63–5.

8. Muñoz LS, Parra B, Pardo CA, Study NEitA. Neurological implications of Zika virus infection in adults. The Journal of infectious diseases. 2017;216(Suppl_10):S897–S905.

9. Platt DJ, Smith AM, Arora N, Diamond MS, Coyne CB, Miner JJ. Zika virus–related neurotropic flaviviruses infect human placental explants and cause fetal demise in mice. Science translational medicine. 2018;10(426):eaao7090.

10. Da Silva IRF, Frontera JA, De Filippis AMB, Do Nascimento OJM, Group R-G-ZR. Neurologic complications associated with the Zika virus in Brazilian adults. JAMA neurology. 2017;74(10):1190–8.

11. Rozé B, Najioullah F, Fergé J-L, Apetse K, Brouste Y, Cesaire R, et al. Zika virus detection in urine from patients with Guillain-Barre syndrome on Martinique, January 2016. Eurosurveillance. 2016;21(9):30154.

12. Azevedo RS, Araujo MT, Martins Filho AJ, Oliveira CS, Nunes BT, Cruz AC, et al. Zika virus epidemic in Brazil. I. Fatal disease in adults: clinical and laboratorial aspects. Journal of Clinical Virology. 2016;85:56–64.

13. Alves-Leon SV, Lima MDR, Nunes PCG, Chimelli LMC, Rabelo K, Nogueira RMR, et al. Zika virus found in brain tissue of a multiple sclerosis patient undergoing an acute disseminated encephalomyelitis-like episode. Multiple Sclerosis Journal. 2019;25(3):427–30.

14. Tang H, Hammack C, Ogden SC, Wen Z, Qian X, Li Y, et al. Zika virus infects human cortical neural progenitors and attenuates their growth. Cell stem cell. 2016;18(5):587–90.

15. Devhare P, Meyer K, Steele R, Ray RB, Ray R. Zika virus infection dysregulates human neural stem cell growth and inhibits differentiation into neuroprogenitor cells. Cell death & disease. 2017;8(10):e3106–e.

16. Li C, Xu D, Ye Q, Hong S, Jiang Y, Liu X, et al. Zika virus disrupts neural progenitor development and leads to microcephaly in mice. Cell stem cell. 2016;19(1):120–6.

17. Quicke KM, Bowen JR, Johnson EL, McDonald CE, Ma H, O’Neal JT, et al. Zika virus infects human placental macrophages. Cell host & microbe. 2016;20(1):83–90.

18. Meertens L, Labeau A, Dejarnac O, Cipriani S, Sinigaglia L, Bonnet-Madin L, et al. Axl mediates ZIKA virus entry in human glial cells and modulates innate immune responses. Cell reports. 2017;18(2):324–33.

19. Lum F-M, Low DK, Fan Y, Tan JJ, Lee B, Chan JK, et al. Zika virus infects human fetal brain microglia and induces inflammation. Clinical Infectious Diseases. 2017;64(7):914–20.

20. Cervellati C, Trentini A, Pecorelli A, Valacchi G. Inflammation in neurological disorders: the thin boundary between brain and periphery. Antioxidants & Redox Signaling. 2020;33(3):191–210.

21. Simi A, Tsakiri N, Wang P, Rothwell N. Interleukin-1 and inflammatory neurodegeneration. Biochemical Society Transactions. 2007;35(5):1122–6.

22. Zhao M-L, Kim M-O, Morgello S, Lee SC. Expression of inducible nitric oxide synthase, interleukin-1 and caspase-1 in HIV-1 encephalitis. Journal of neuroimmunology. 2001;115(1-2):182–91.

23. Xing HQ, Hayakawa H, Izumo K, Kubota R, Gelpi E, Budka H, et al. In vivo expression of proinflammatory cytokines in HIV encephalitis: an analysis of 11 autopsy cases. Neuropathology. 2009;29(4):433–42.

24. Bonifati DM, Kishore U. Role of complement in neurodegeneration and neuroinflammation. Molecular immunology. 2007;44(5):999–1010.

25. Barnum SR, Jones JL, Benveniste EN. Interferon-gamma regulation of C3 gene expression in human astroglioma cells. Journal of neuroimmunology. 1992;38(3):275–82.

26. Barnum SR, Jones JL, Benveniste EN. Interleukin-1 and tumor necrosis factor-mediated regulation of C3 gene expression in human astroglioma cells. Glia. 1993;7(3):225–36.

27. Cardinaux JR, Allaman I, Magistretti PJ. Pro-inflammatory cytokines induce the transcription factors C/EBPβ and C/EBPδ in astrocytes. Glia. 2000;29(1):91–7.

28. Speth C, Williams K, Hagleitner M, Westmoreland S, Rambach G, Mohsenipour I, et al. Complement synthesis and activation in the brain of SIV-infected monkeys. Journal of neuroimmunology. 2004;151(1-2):45–54.

29. Basu A, Krady JK, Levison SW. Interleukin-1: a master regulator of neuroinflammation. Journal of neuroscience research. 2004;78(2):151–6.

30. Grant A, Ponia SS, Tripathi S, Balasubramaniam V, Miorin L, Sourisseau M, et al. Zika virus targets human STAT2 to inhibit type I interferon signaling. Cell host & microbe. 2016;19(6):882–90.

31. Kumar A, Hou S, Airo AM, Limonta D, Mancinelli V, Branton W, et al. Zika virus inhibits type-I interferon production and downstream signaling. EMBO reports. 2016;17(12):1766–75.

32. Lazear HM, Govero J, Smith AM, Platt DJ, Fernandez E, Miner JJ, et al. A mouse model of Zika virus pathogenesis. Cell host & microbe. 2016;19(5):720–30.

33. Bowen JR, Quicke KM, Maddur MS, O’Neal JT, McDonald CE, Fedorova NB, et al. Zika virus antagonizes type I interferon responses during infection of human dendritic cells. PLoS pathogens. 2017;13(2):e1006164.

34. Jurado KA, Yockey LJ, Wong PW, Lee S, Huttner AJ, Iwasaki A. Antiviral CD8 T cells induce Zika-virus-associated paralysis in mice. Nature microbiology. 2018;3(2):141–7.

35. Morrey JD, Oliveira AL, Wang H, Zukor K, de Castro MV, Siddharthan V. Zika virus infection causes temporary paralysis in adult mice with motor neuron synaptic retraction and evidence for proximal peripheral neuropathy. Scientific reports. 2019;9(1):1–15.

36. Jeong GU, Lyu J, Kim K-D, Chung YC, Yoon GY, Lee S, et al. SARS-CoV-2 Infection of Microglia Elicits Proinflammatory Activation and Apoptotic Cell Death. Microbiology Spectrum. 2022:e01091–22.

37. Xu P, Shan C, Dunn TJ, Xie X, Xia H, Gao J, et al. Role of microglia in the dissemination of Zika virus from mother to fetal brain. PLoS neglected tropical diseases. 2020;14(7):e0008413.

38. Richard AS, Shim B-S, Kwon Y-C, Zhang R, Otsuka Y, Schmitt K, et al. AXL-dependent infection of human fetal endothelial cells distinguishes Zika virus from other pathogenic flaviviruses. Proceedings of the National Academy of Sciences. 2017;114(8):2024–9.

39. Yang D, Chu H, Lu G, Shuai H, Wang Y, Hou Y, et al. STAT2-dependent restriction of Zika virus by human macrophages but not dendritic cells. Emerging microbes & infections. 2021;10(1):1024–37.

40. Faria SS, Costantini S, de Lima VCC, de Andrade VP, Rialland M, Cedric R, et al. NLRP3 inflammasome-mediated cytokine production and pyroptosis cell death in breast cancer. Journal of Biomedical Science. 2021;28(1):1–15.

41. Hernandez-Encinas E, Aguilar-Morante D, Cortes-Canteli M, Morales-Garcia JA, Gine E, Santos A, et al. CCAAT/enhancer binding protein β directly regulates the expression of the complement component 3 gene in neural cells: implications for the pro-inflammatory effects of this transcription factor. Journal of neuroinflammation. 2015;12(1):1–16.

42. Mazumdar B, Kim H, Meyer K, Bose SK, Di Bisceglie AM, Ray RB, et al. Hepatitis C virus proteins inhibit C3 complement production. Journal of virology. 2012;86(4):2221–8.

43. Yaron M, ShirazI I, Yaron I. eAnti-interleukin-1 effects of diacerein and rhein in human osteoarthritic synovial tissue and cartilage cultures. Osteoarthritis and Cartilage. 1999;7(3):272–80.

44. Klein RS, Garber C, Funk KE, Salimi H, Soung A, Kanmogne M, et al. Neuroinflammation during RNA viral infections. Annual review of immunology. 2019;37:73.

45. Figueiredo CP, Barros-Aragão FG, Neris RL, Frost PS, Soares C, Souza IN, et al. Zika virus replicates in adult human brain tissue and impairs synapses and memory in mice. Nature communications. 2019;10(1):1–16.

46. Kalia M, Khasa R, Sharma M, Nain M, Vrati S. Japanese encephalitis virus infects neuronal cells through a clathrin-independent endocytic mechanism. Journal of virology. 2013;87(1):148–62.

47. Klein RS, Lin E, Zhang B, Luster AD, Tollett J, Samuel MA, et al. Neuronal CXCL10 directs CD8+ T-cell recruitment and control of West Nile virus encephalitis. Journal of virology. 2005;79(17):11457–66.

48. Delhaye S, Paul S, Blakqori G, Minet M, Weber F, Staeheli P, et al. Neurons produce type I interferon during viral encephalitis. Proceedings of the National Academy of Sciences. 2006;103(20):7835–40.

49. Chevalier G, Suberbielle E, Monnet C, Duplan V, Martin-Blondel G, Farrugia F, et al. Neurons are MHC class I-dependent targets for CD8 T cells upon neurotropic viral infection. PLoS pathogens. 2011;7(11):e1002393.

50. Ginhoux F, Greter M, Leboeuf M, Nandi S, See P, Gokhan S, et al. Fate mapping analysis reveals that adult microglia derive from primitive macrophages. Science. 2010;330(6005):841–5.

51. Xiong X-Y, Liu L, Yang Q-W. Functions and mechanisms of microglia/macrophages in neuroinflammation and neurogenesis after stroke. Progress in neurobiology. 2016;142:23–44.

52. Yamasaki R, Lu H, Butovsky O, Ohno N, Rietsch AM, Cialic R, et al. Differential roles of microglia and monocytes in the inflamed central nervous system. Journal of Experimental Medicine. 2014;211(8):1533–49.

53. Hafer-Macko C, Sheikh K, Li C, Ho T, Cornblath D, McKhann G, et al. Immune attack on the Schwann cell surface in acute inflammatory demyelinating polyneuropathy. Annals of Neurology: Official Journal of the American Neurological Association and the Child Neurology Society. 1996;39(5):625–35.

54. Dinarello CA. Interleukin-1 in the pathogenesis and treatment of inflammatory diseases. Blood, The Journal of the American Society of Hematology. 2011;117(14):3720–32.

55. McKenzie BA, Dixit VM, Power C. Fiery cell death: pyroptosis in the central nervous system. Trends in neurosciences. 2020;43(1):55–73.

56. Ramaglia V, Daha M, Baas F. The complement system in the peripheral nerve: friend or foe? Molecular immunology. 2008;45(15):3865–77.

57. Guo R-F, Ward PA. Role of C5a in inflammatory responses. Annu Rev Immunol. 2005;23:821–52.

58. Kuroda H, Fujihara K, Takano R, Takai Y, Takahashi T, Misu T, et al. Increase of complement fragment C5a in cerebrospinal fluid during exacerbation of neuromyelitis optica. Journal of neuroimmunology. 2013;254(1-2):178–82.

59. Giorgio C, Zippoli M, Cocchiaro P, Castelli V, Varrassi G, Aramini A, et al. Emerging role of C5 complement pathway in peripheral neuropathies: Current treatments and future perspectives. Biomedicines. 2021;9(4):399.

60. Shim B-S, Kwon Y-C, Ricciardi MJ, Stone M, Otsuka Y, Berri F, et al. Zika virus-immune plasmas from symptomatic and asymptomatic individuals enhance Zika pathogenesis in adult and pregnant mice. MBio. 2019;10(4):e00758–19.

61. Sanjana NE, Shalem O, Zhang F. Improved vectors and genome-wide libraries for CRISPR screening. Nature methods. 2014;11(8):783–4.

62. Stringer BW, Day BW, D’Souza RC, Jamieson PR, Ensbey KS, Bruce ZC, et al. A reference collection of patient-derived cell line and xenograft models of proneural, classical and mesenchymal glioblastoma. Scientific reports. 2019;9(1):1–14.

63. Cardona AE, Huang D, Sasse ME, Ransohoff RM. Isolation of murine microglial cells for RNA analysis or flow cytometry. Nature protocols. 2006;1(4):1947–51.

64. Garcia JA, Cardona SM, Cardona AE. Isolation and analysis of mouse microglial cells. Current protocols in immunology. 2014;104(1):14.35.1-14.35.15.

65. Martin M. Cutadapt removes adapter sequences from high-throughput sequencing reads. EMBnet journal. 2011;17(1):10–2.

66. Dobin A, Davis CA, Schlesinger F, Drenkow J, Zaleski C, Jha S, et al. STAR: ultrafast universal RNA-seq aligner. Bioinformatics. 2013;29(1):15–21.

67. Anders S, Pyl PT, Huber W. HTSeq—a Python framework to work with high-throughput sequencing data. bioinformatics. 2015;31(2):166–9.

68. Love MI, Huber W, Anders S. Moderated estimation of fold change and dispersion for RNA-seq data with DESeq2. Genome biology. 2014;15(12):1–21.

69. Kuleshov MV, Jones MR, Rouillard AD, Fernandez NF, Duan Q, Wang Z, et al. Enrichr: a comprehensive gene set enrichment analysis web server 2016 update. Nucleic acids research. 2016;44(W1):W90–W7.

